# A comprehensive evaluation of consensus spectrum generation methods in proteomics

**DOI:** 10.1101/2022.01.25.477699

**Authors:** Xiyang Luo, Wout Bittremieux, Johannes Griss, Eric W Deutsch, Timo Sachsenberg, Lev I. Levitsky, Mark V. Ivanov, Julia A. Bubis, Ralf Gabriels, Henry Webel, Aniel Sanchez, Mingze Bai, Lukas Kall, Yasset Perez-Riverol

## Abstract

Spectrum clustering is a powerful strategy to minimize redundant mass spectral data by grouping highly similar mass spectra corresponding to repeatedly measured analytes. Based on spectrum similarity, near-identical spectra are grouped in clusters, after which each cluster can be represented by its so-called consensus spectrum for downstream processing. Although several algorithms for spectrum clustering have been adequately benchmarked and tested, the influence of the consensus spectrum generation step is rarely evaluated. Here, we present an implementation and benchmark of common consensus spectrum algorithms, including spectrum averaging, spectrum binning, the most similar spectrum, and the best-identified spectrum. We have analyzed diverse public datasets using two different clustering algorithms (spectra-cluster and MaRaCluster) to evaluate how the consensus spectrum generation procedure influences downstream peptide identification. The BEST and BIN methods were found the most reliable methods for consensus spectrum generation, including for datasets with post-translational modifications (PTM) such as phosphorylation. All source code and data of the present study are freely available on GitHub at https://github.com/statisticalbiotechnology/representative-spectra-benchmark.

## Introduction

Spectrum clustering, i.e. the process of grouping similar spectra in a larger collection of MS2 spectra into smaller subsets, has multiple applications in mass spectrometry in general and in proteomics in particular (1), including the generation of spectral libraries (2) and spectral archives (3), quality assessment of peptide identifications in public repositories (2) and improvement of quantification results (4, 5). Spectrum clustering algorithms strive to group highly similar spectra so that each cluster contains spectra generated from the same analyte (peptidoforms with a specific charge in the case of proteomics). Differences between tools for spectrum clustering vary in their implementation of the various data processing steps, including the pre-processing of spectra (e.g., intensity normalization and peak picking), the clustering algorithm used, the metric used for determining similarity between spectra, and the optional optimizations to increase computational efficiency. Current tools for spectrum clustering include MS-Cluster (3), spectra-cluster (2), MaRaCluster (6), msCRUSH (7), and falcon (8).

While the most apparent output of the process of spectrum clustering is a grouping of spectra into clusters, the majority of use cases benefit from a condensed single spectrum representation for each cluster. This is, for instance, useful for the data-driven creation of spectral libraries (9), for reannotation and visualization of clustering results in public data repositories (2), and label-free quantification (4). The generation of high-quality representative spectra for each cluster is a key aspect of spectrum clustering, as the resulting consensus spectra form the starting point for downstream analyses. Although several spectrum clustering algorithms have been adequately benchmarked [8], the impact of the consensus spectrum generation procedure has so far not been properly evaluated. Several common approaches can be used to generate representative spectra, including spectrum binning, spectrum averaging (3), and selecting the most similar spectrum to all cluster members (medoid) (8). Additionally, although this strategy can only be used for clusters that contain one or more identified spectra, the “best-identified spectrum” method uses the most confidently identified spectrum as cluster representative (10, 11)

Here, we have performed a comprehensive evaluation of algorithms for the generation of consensus spectra to assess their performance for downstream processing of spectrum clustering results. We have used the spectra-cluster and MaRaCluster tools to generate clusters from diverse publicly available datasets and explore whether consensus spectrum generation algorithms perform differently between different tools. Additionally, we have evaluated the impact of consensus spectrum generation on downstream peptide and protein identification performance. All code and analyses are open-source and available at https://github.com/statisticalbiotechnology/representative-spectra-benchmark under the permissive Apache 2.0 license.

## Methods

### Consensus spectrum generation algorithms and evaluation

For the benchmark, we implemented four consensus spectrum generation algorithms:

- Spectrum averaging (AVERAGE): The representative spectrum is an average of all the spectra in the cluster (9, 12, 13). In this algorithm, for every m/z value, the corresponding intensities on each spectrum in the cluster are averaged.
- Spectrum binning (BIN): In this method, for each cluster, a consensus spectrum vector with bin width 0.02 *m*/*z* was first constructed (13). For all spectra in the cluster, peak m/z and intensity values were assigned to the corresponding bin in the consensus spectrum vector. Bins that contained values from fewer than 25% of the cluster members were discarded. Next, the vector was converted to a consensus spectrum by averaging all peak m/z and intensity values per bin (2).
- Most similar spectrum (MOST): For each cluster, the spectrum that is on average most similar to all cluster members was selected as representative (14, 15). This was determined by first calculating the dot product of all pairwise similarities between spectra in the cluster. Next, the spectrum with the maximal summed dot product to all other spectra was selected as the representative for that cluster.
- Best identified spectrum (BEST): For each cluster that contained at least one identified spectrum, the spectrum with the maximal peptide-spectrum match score was chosen as the representative for that cluster. Note that this approach is not valid if all spectra in the cluster are unmatched.

Data manipulation steps were implemented as reproducible Nextflow workflows (**Figure 1**). The spectra-cluster (version 1.1.2) (2) and MaRaCluster (version 1.0) (6) spectrum clustering tools were used to cluster the mass spectrum data, and the MS-GF+ sequence database search engine (version v2021.03.22) (16) was used to perform peptide identification. For each cluster, representative (consensus) spectra were directly generated from the clustering output using the first three consensus generation procedures described above. For the best-identified method, the spectra were additionally identified using MS-GF+, after which the PSMs with the maximum scores were selected as representatives for each cluster. To ensure a fair comparison between all consensus spectrum generation procedures, clusters that only contained unidentified spectra were ignored, as no valid representative spectrum could be obtained using the best-identified method. To evaluate downstream peptide identification performance, the consensus spectra obtained for both spectrum clustering tools with each consensus spectrum generation method were searched using MS-GF+ (16), after which the number of peptide identifications was compared between all combinations of clustering and consensus generation methods, and with the original data without clustering.

**Figure 1:**
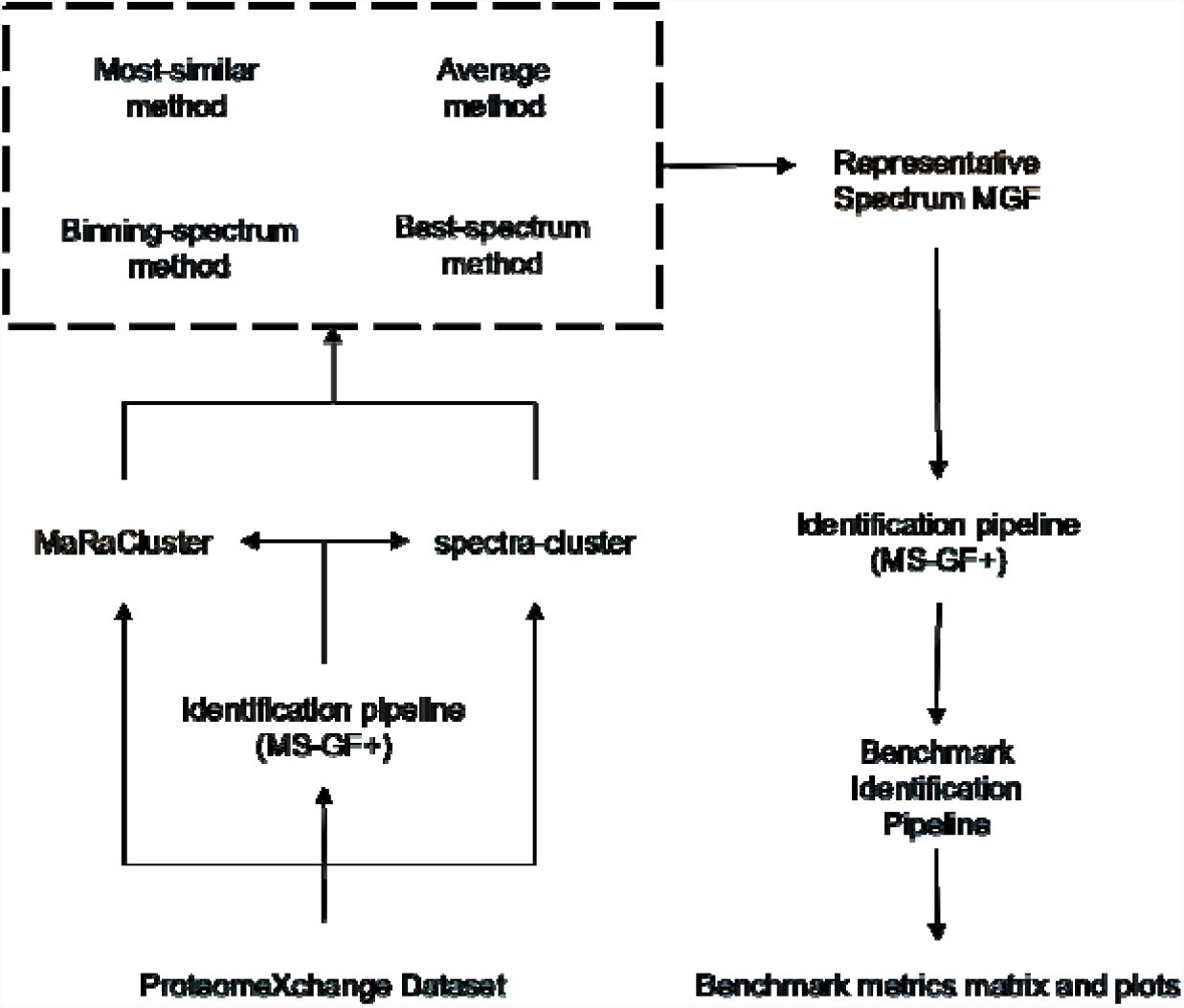
Study workflow, including clustering and peptide identification of publicly available ProteomeXchange datasets, consensus spectrum generation using alternative procedures, and evaluation of cluster representatives using an identification benchmark.

### Benchmark Datasets

We used four public ProteomeXchange datasets: PXD008355, PXD023047, PXD021518, and PXD023361 (**Table 1**). RAW data from each dataset was converted to MGF using the ThermoRawFileParser (version: 1.2.3) tool (17) with default parameters. Among them, PXD008355, PXD023047, and PXD021518 are from Arabidopsis thaliana (mouse-ear cress), and PXD023361 is from Saccharomyces cerevisiae (baker’s yeast). The datasets have been acquired using three different instrument models: Q Exactive, Q Exactive HF, and Q Exactive HF-X. The description of the samples, instrument configuration, sample processing steps, and analytical method can be read in the original publications: PXD008355 (18), PXD023047 (19), PXD021518 (20), and PXD023361 (21).

**Table 1.** Datasets were reanalysed to evaluate the performance of each consensus spectrum generation algorithm. The number of peptide identifications and peptide-spectrum matches can be found in the Supplementary Notes. In addition, the description of each dataset can be found in the original publication and PRIDE Archive (22).

| Project accession | Instrument | No. MS/MS |
| --- | --- | --- |
| PXD008355 (18) | Q Exactive | 1,477,567 |
| PXD023047 (19) | Q Exactive HF | 109,333 |
| PXD021518 (20) | Q Exactive HF-X | 286,410 |
| PXD023361 (21) | Q Exactive | 38,286 |

For datasets PXD008355, PXD023047, and PXD021518, the Arabidopsis Thaliana protein database was downloaded from http://ftp.ebi.ac.uk/pride-archive/2019/07/PXD008355/TAIR10.fasta, while for dataset PXD023361 the Saccharomyces cerevisiae database was downloaded from http://ftp.pride.ebi.ac.uk/pride/data/archive/2021/04/PXD023361/uniprot-S_yeast.fasta.

For datasets PXD008355, PXD021518, and PXD023361, the precursor error tolerance was set to 10 ppm; while for dataset PXD023047, it was set to 20ppm. Target-decoy was performed using MS-GF+ (parameter -tda). For datasets PXD023047 and PXD021518 two modifications were allowed (NumMods=2): fixed carbamidomethyl cysteine modification, and variable methionine oxidation; while for datasets PXD008355 and PXD023361 Phosphorylation was also considered as variable modification.

### Code availability

All code and analyses are freely available as open source under the Apache 2.0 license at https://github.com/statisticalbiotechnology/representative-spectra-benchmark. The consensus generation procedures were implemented in Python 3.6. Software dependencies that were used include Matplotlib (version 3.1.2) (23), Numba (version 0.47.0) (24), NumPy (version 1.17.3) (25), Pandas (version 0.25.3), pyOpenMS (version 2.4.0) (26), Pyteomics (version 4.1.2) (27), and spectrum_utils (version 0.3.3) (28).

## Results

Figure 2 shows the number of PSMs (FDR=1%) identified with MS-GF+ (datasets PXD023047, PXD021528, PXD008355, and PXD023361) for spectrum clustering using MaRaCluster and spectra-cluster followed by consensus spectrum generation using the MOST, AVERAGE, BIN, and BEST procedures. Among the four public proteomics datasets, whether using spectrum clustering results from MaRaCluster or spectra-cluster, the identification rate for the MOST method is lower compared to the other methods, while the BIN and BEST methods achieve a higher spectrum identification rate (**Figure 2**).

**Figure 2:**
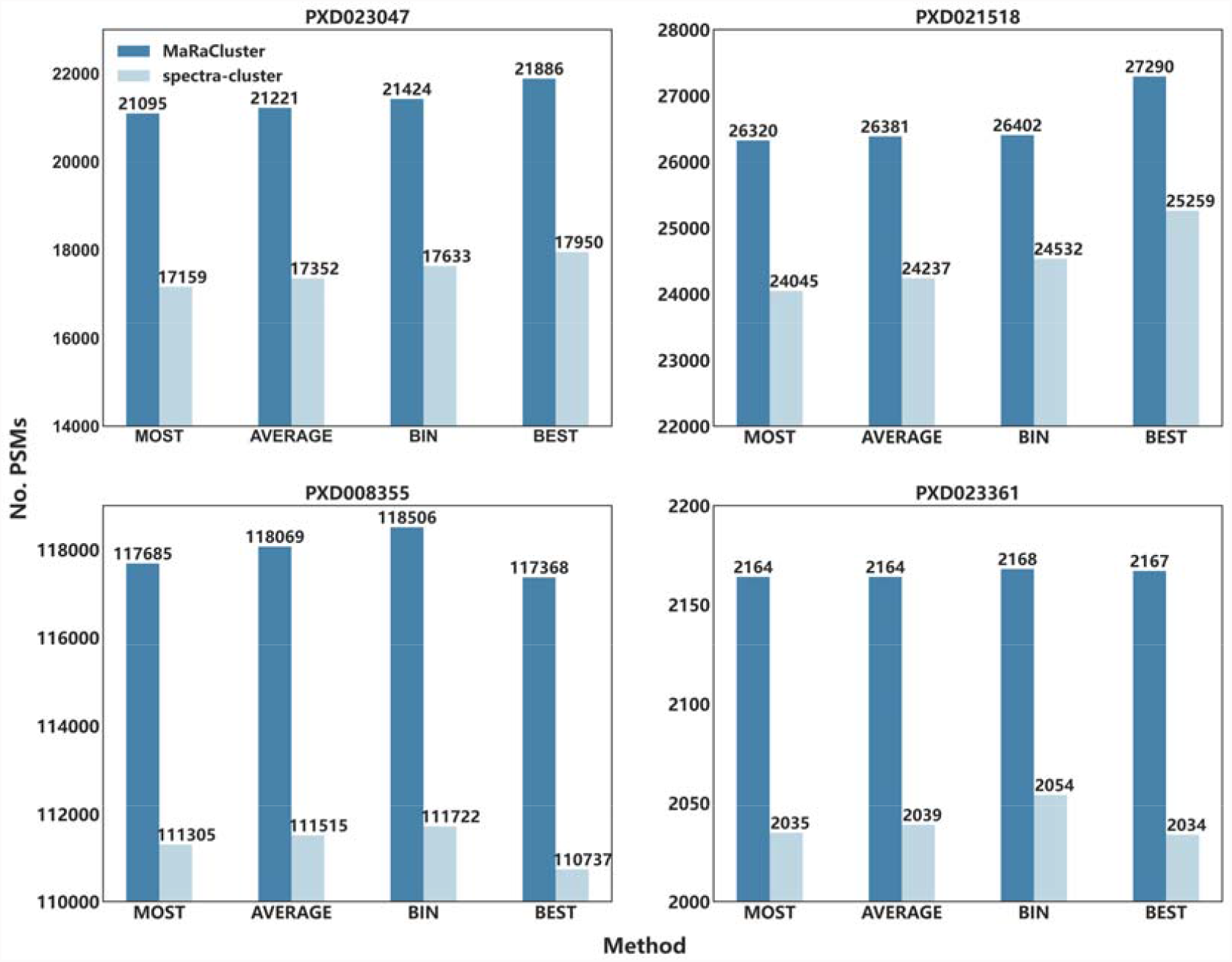
The number of PSMs obtained by MS-GF+ when searching consensus spectra produced by the MOST, AVERAGE, BIN, and BEST representative cluster generation methods for public proteomics datasets PXD023047, PXD021528, PXD008355, and PXD023361. Note that the bar plots are truncated past 0 to highlight relevant performance differences.

While the number of identified spectra only differs by a small amount between the various consensus spectrum generation procedures, when analyzing big public proteomics databases (billions of spectra) (29) these differences can be translated into millions of spectrum identifications. Among the methods that transform the original spectra, the BIN method is the one that performs best. The BIN method divides the m/z values into small bins and then overlaps multiple spectra within those bins. If there are multiple intensities in a bin, the algorithm will superimpose intensities in the same bin, favoring the most intensive peaks, which could improve peptide identification. However, in some cases, can also remove important peaks from the MS/MS spectra.

Most of the consensus generation methods modify the original spectra, not only by removing or keeping some of the spectrum peaks, but also by modifying the corresponding intensity of each peak. We have used the distributions of the MS-GF+ RawScore to explore the relationship between the final spectra and the quality of the peptide identifications. Figure 3 shows the distribution of MS-GF+ RawScore for the four consensus generation methods (MOST, AVERAGE, BIN, and BEST) after clustering with MaRaCluster and spectra-cluster. For both clustering tools, the BIN and BEST method generate consensus spectra with higher average RawScore values (**Figure 3**), and similar to the previous metric (number of PSMs), the BEST algorithm achieves the highest average RawScore (**Supplementary Note 1**). The representative consensus spectra generated by the MOST method have the lowest average RawScore (**Figure 3**). The distribution of RawScore values (**Figure 3**) shows that the RawScores are more homogenous for the BIN method (lower standard deviation) than for all the other methods, including the BEST algorithm (**Supplementary Note 1**).

**Figure 3:**
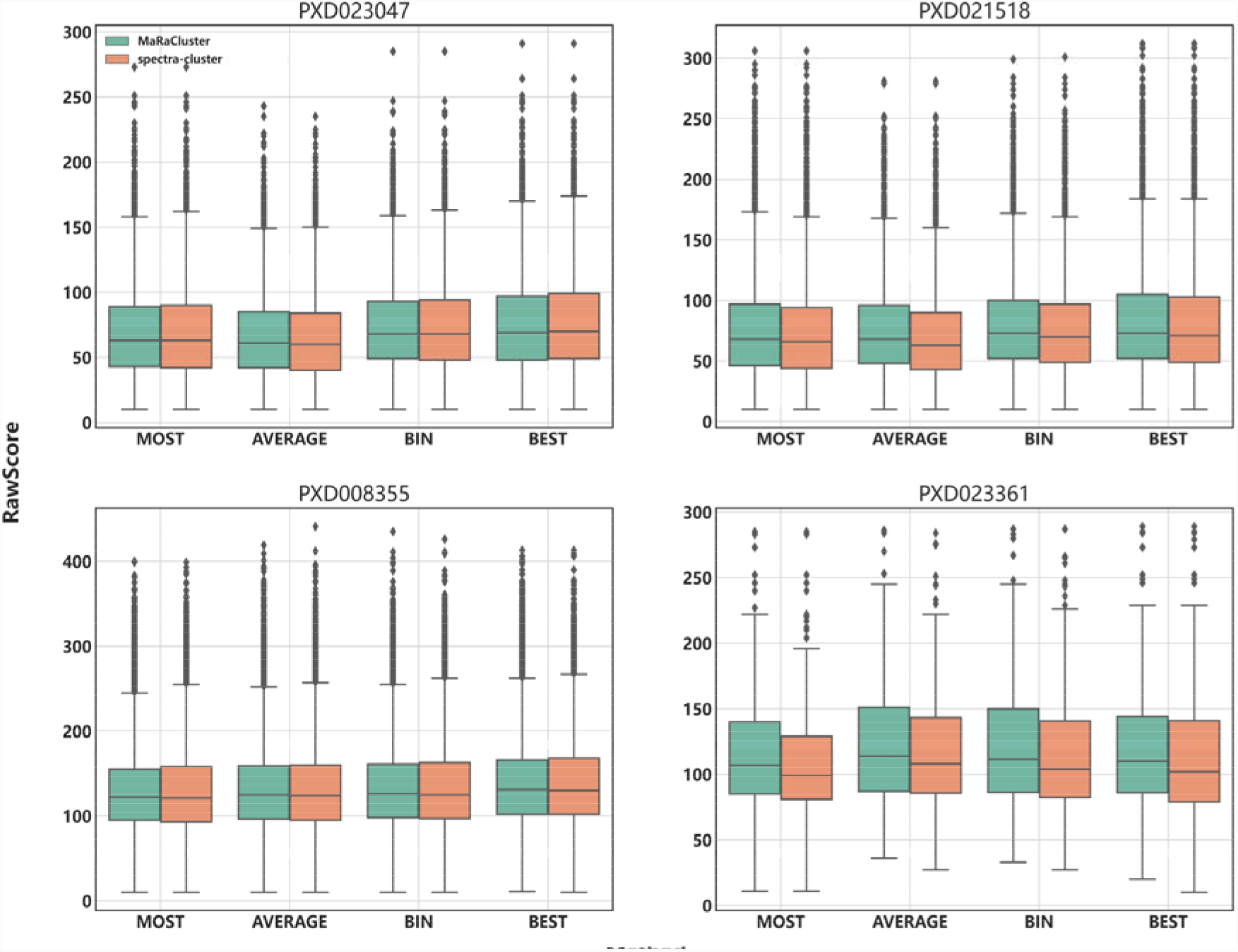
Distribution of MS-GF+ RawScores for MOST, AVERAGE, BIN, and BEST representative spectra from the public proteomics datasets PXD023047, PXD021528, PXD008355, and PXD023361.

Figure 4 shows the changes in mean RawScore of the identified spectra generated with the four evaluated methods (MOST, AVERAGE, BIN, and BEST) for clusters of different sizes (cluster sizes 1, 2, 3, 4, 5, 5-10, 10-20, 20 or higher). As expected, for clusters of one spectrum, no differences have been seen between different consensus methods, but minor differences are observed between clustering algorithms. For other small clusters containing three or fewer spectra, consensus spectra derived from the spectra-cluster results, in combinations with all the consensus spectrum generation methods, provide higher mean RawScores than consensus spectra derived from MaRaCluster results. In contrast, for larger clusters, MaRaCluster consensus spectra lead to higher mean RawScores. For both spectra-cluster and MaRaCluster, the mean RawScore increases with increasing cluster size. The BEST and BIN algorithms are stable for both clustering algorithms and all datasets (**Supplementary Note 1**), and the scores of these two algorithms are generally higher than MOST and AVERAGE. In combination with MaRaCluster, the AVERAGE algorithm shows instability and the score of the AVERAGE algorithm is generally lower than the other three algorithms.

**Figure 4:**
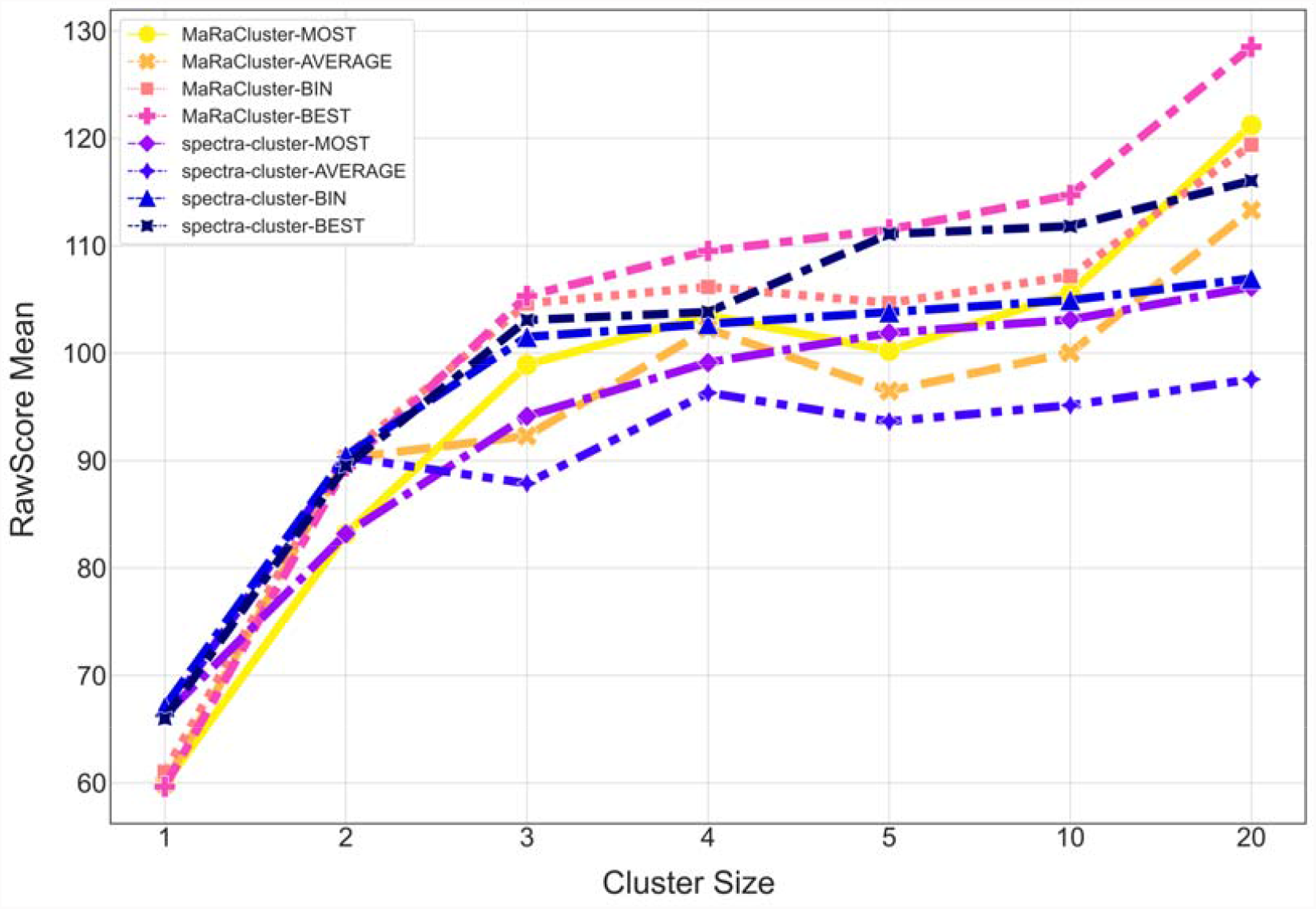
RawScore mean of the four different evaluated methods under different cluster sizes (1, 2, 3, 4, 5, 5-10, 10-20, 20 or higher).

In addition to peptide identification, we explored how using consensus spectra instead of the original spectra affects phospho-peptide identification and phosphorylation site localization.

We analyzed the number of phosphorylation sites identified in dataset PXD008355 after clustering with both tools (MaRaCluster and spectra-cluster) and the four different consensus spectrum generation methods (MOST, AVERAGE, BIN, and BEST). We have evaluated two metrics, (i) the number of phosphorylated PSMs identified and (ii) the phosphorylation sites identified.

Figure 5 shows the intersection of the phosphorylated PSMs among the four representative cluster methods after spectrum clustering with MaRaCluster and spectra-cluster. Most of the PSMs (91.2% for MaRaCluster and 96.4% for spectra-cluster) for the four representative cluster methods produce the same phosphorylated PSMs. The BIN method produces the largest number of unique PSMs, which is about double the number of other methods, followed by the BEST, MOST, and AVERAGE methods (**Figure 5**).

**Figure 5:**
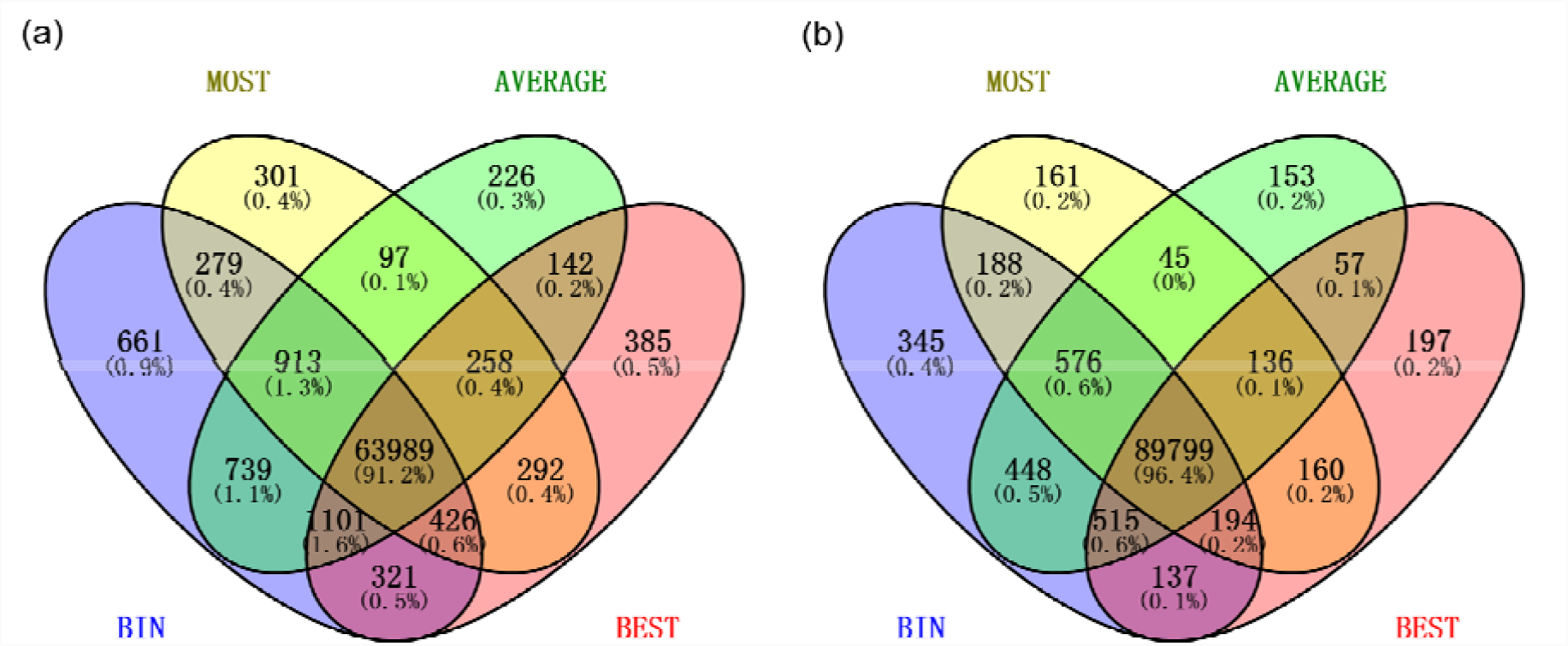
The intersection of the total phosphorylated PSMs among the four representative cluster methods in **(a)** MaRaCluster, and **(b)** spectra-cluster.

While the majority of phosphorylated PSMs are aggregated among all methods, around ∼1% are different and we also observed differences in terms of phosphorylation sites. Table 2 shows the difference in phosphorylation sites between BIN and BEST representative spectra from MaRaCluster and spectra-cluster (extended table, **Supplementary Notes 3**). Because the BEST and BIN methods were the best performing consensus generation (13) options in terms of peptide identification, we focus the discussion on these two methods (extended table, Supplementary Notes 3). Most phosphorylated PSMs (63,165 for MaRaCluster and 89,161 for spectra-cluster) for the BEST and BIN methods are identical. However, a small number of phospho sites are different (2683 in MaRaCluster and 1494 in spectra-cluster), some of them due to different peptide identifications, some of them due to differences in the localization accuracy after clustering and the change of the spectra. These small differences can be attributed to the fact that the BIN method modifies the ion peak intensity and m/z of the spectrum through the binning algorithm.

**Table 2:**
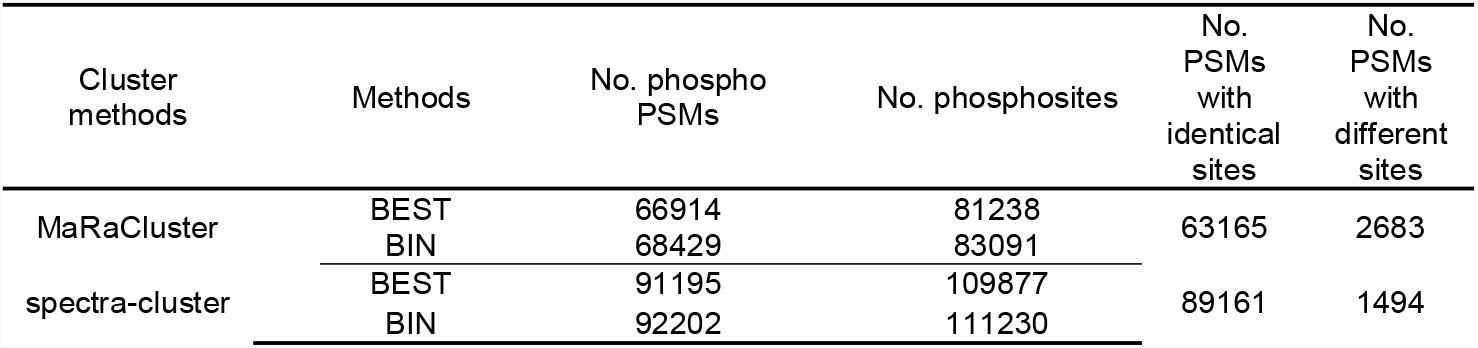
Analysis of phosphorylation sites identification of dataset PXD008355, after clustering with MaRaCluster and spectra-cluster, and generation of the consensus spectra using two different methods (BEST, BIN). We quantified the number of total phosphorylated PSMs and phosphorylation sites for each combination of clustering method and consensus generation method. In addition, we added the number of identical and different phospho-sites between the BEST and BIN methods for each clustering algorithm.

## Conclusions

Representative spectra from clusters have typically been generated using four different algorithms: spectrum averaging, spectrum binning, the most similar spectrum, and the best-identified spectrum. Most tools and resources, including SpectraST (9), MassIVE (11, 30) spectral libraries, or spectra-cluster and PRIDE Cluster (2) use one of these methods. However, to our knowledge, no systematic analysis has been performed to compare multiple algorithms to generate consensus spectra. We implemented a Python framework to benchmark existing algorithms to generate representative spectra from clustering results from two different popular clustering tools—MaRaCluster and spectra-cluster.

The BEST and BIN methods were found to be the most reliable methods for consensus spectrum generation, including for datasets with post-translational modifications such as phosphorylation. The BEST method generates representative consensus spectra based on existing spectrum identification results, which requires that all clusters contain identified spectra. Therefore, the BEST method cannot be used on spectral archives (clusters of non-identified spectra) or if clustering is performed before the identification step. The BIN method is based on the original spectrum file and binning algorithm to generate representative consensus spectra and performed best in all benchmarks and comparisons after the BEST method. While the BIN algorithm modifies the original spectra, we do not observe major differences in identifying phosphorylated peptides and phosphorylation sites compared to the results of the BEST method to generate representative spectra. The fact that the BEST method is performing so well, compared to existing methods, suggests that better algorithms could be developed in the future to generate consensus spectra from clustering results.

## Acknowledgment

The authors would like to acknowledge the EuBIC-MS community that organized the EuBIC-MS Developer Meeting in January 2020 (31), triggering the original discussions and implementations of this work.

