## Supplementary Notes and Figures for "A comprehensive evaluation of consensus spectrum generation methods in proteomics"

### **Supplementary Note 1: Identification of clustering data**

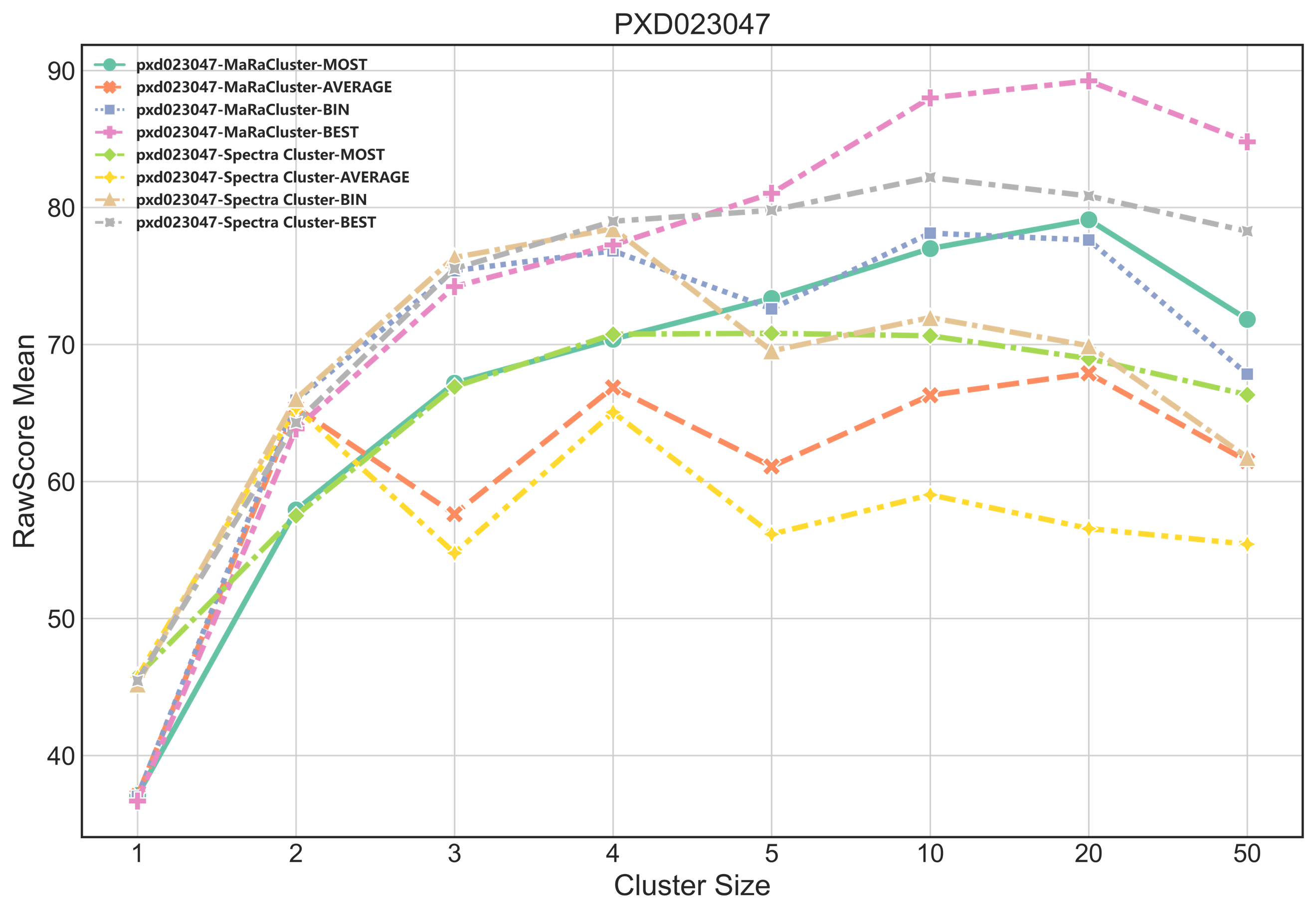

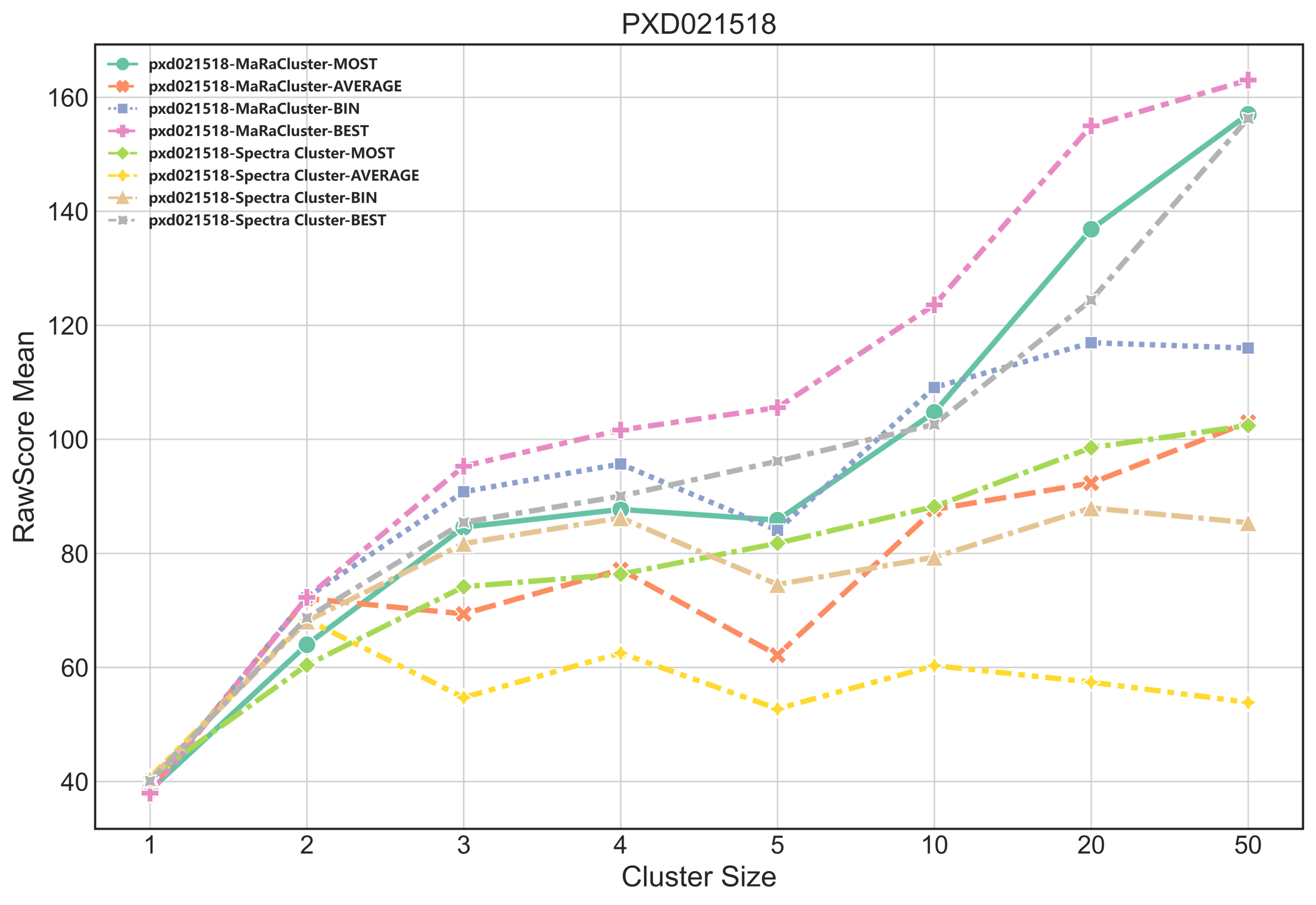

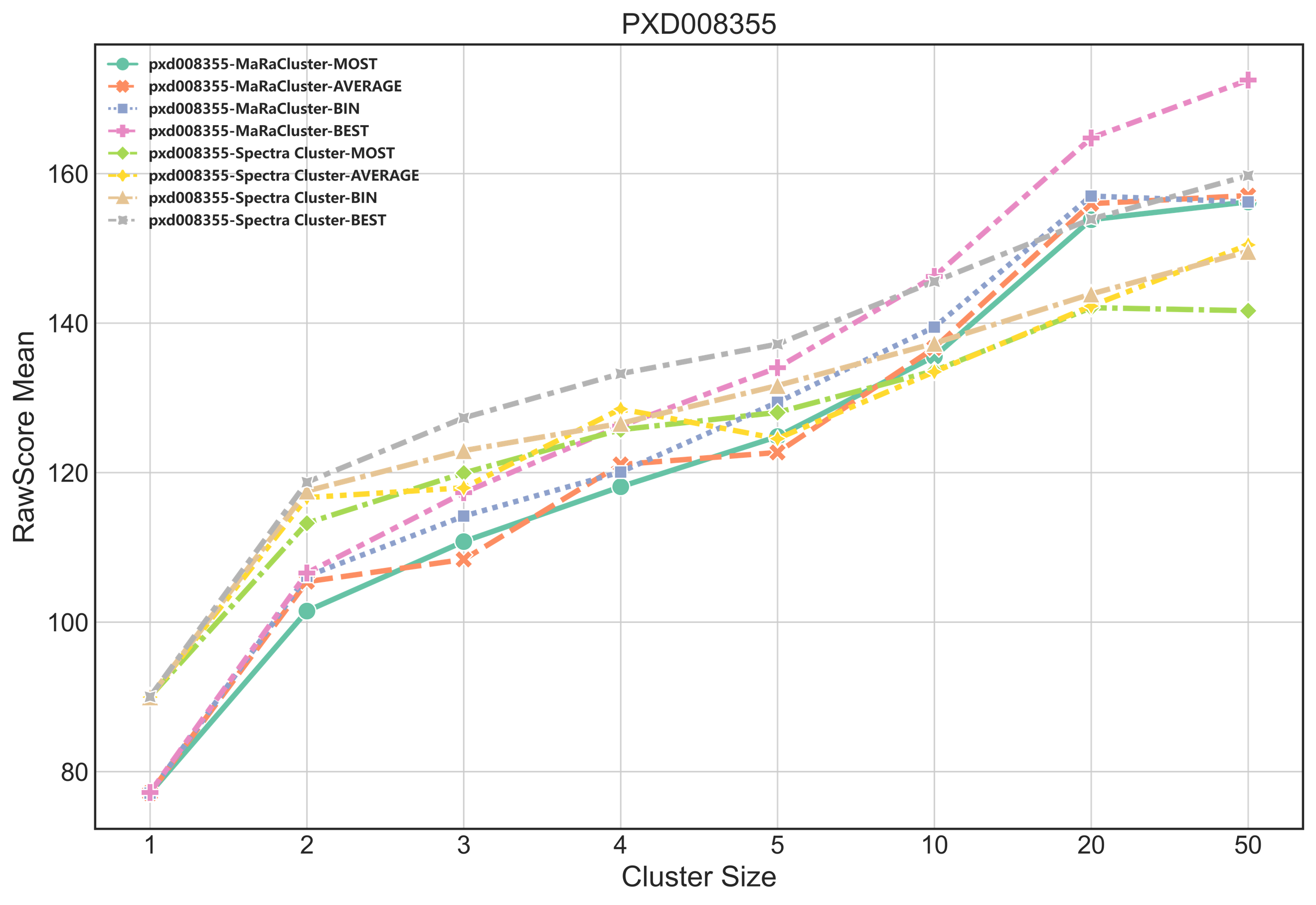

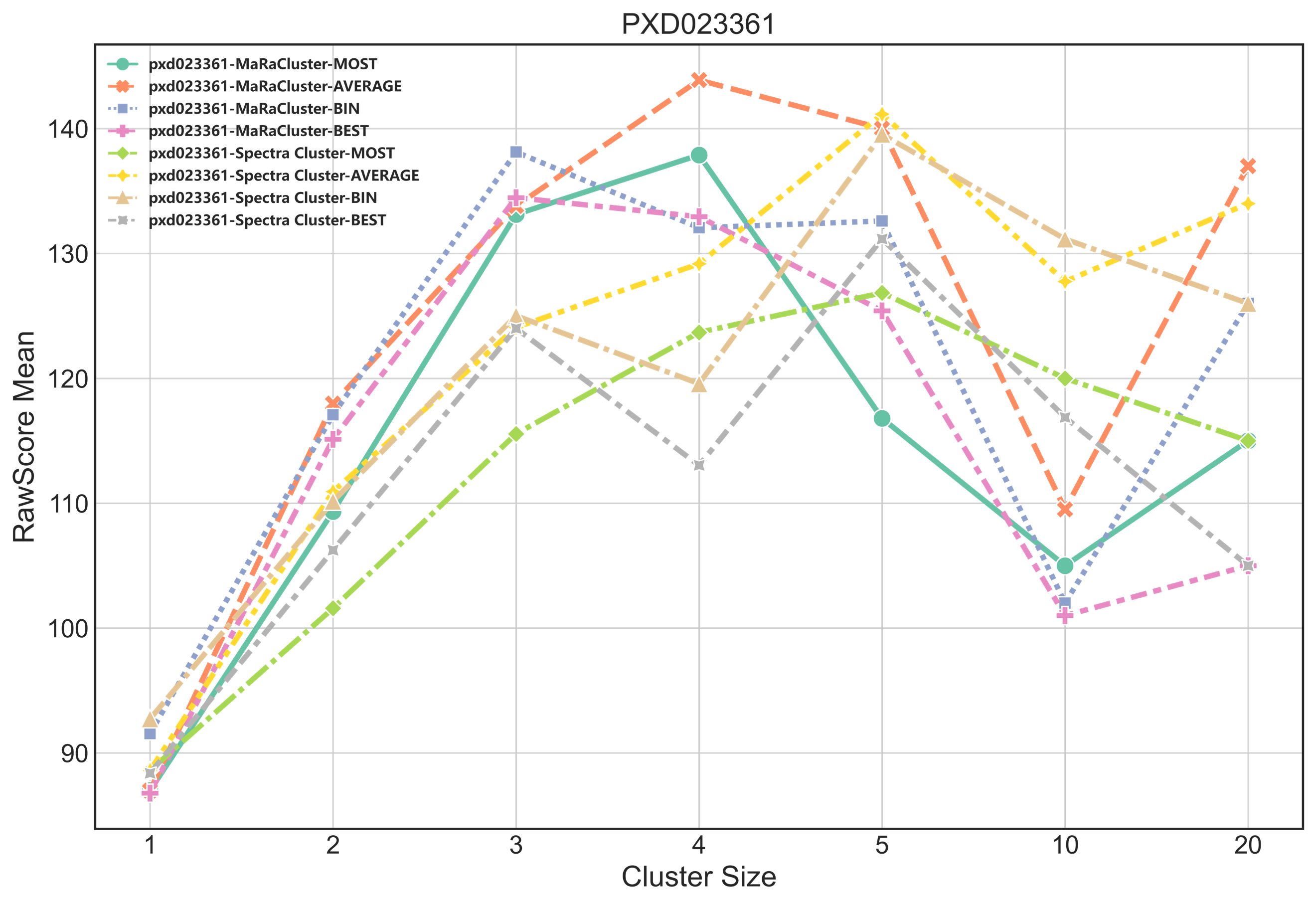

**Figure S3**: RawScore mean changes of the four consistent spectra under different cluster sizes for public proteomics datasets PXD023047, PXD021528, PXD008355 and PXD023361.

### **Supplementary Note 2: The benchmark datasets**

PXD008355

| **MS name** |  | **No. MS** | **No. Unique Peptide** | **No. Unique Peptide**  **（FDR=1）** | **No. PSMs** | **No. PSMs**  **（FDR=1）** | **composite cluster RawScore mean** | **composite cluster Evalue mean** |
| --- | --- | --- | --- | --- | --- | --- | --- | --- |
| Original MS |  | 1477567 | 278562 | 18153 | 772177 | 218794 | - | - |
| MARACluster BEST |  | 945661 | 252903 | 17283 | 540966 | 117368 | 139.77 | 0.00053 |
| MARACluster BIN |  | 1126561 | 250035 | 17355 | 539077 | 118506 | 133.28 | 0.00101 |
| MARACluster MOST |  | 1126561 | 253212 | 17282 | 544324 | 117685 | 128.62 | 0.00115 |
| MARACluster AVERAGE |  | 1126561 | 252316 | 17310 | 544667 | 118069 | 131.03 | 0.00109 |
| spectra-cluster BEST |  | 914502 | 248435 | 17181 | 522299 | 110737 | 144.73 | 0.00053 |
| spectra-cluster BIN |  | 1092343 | 246137 | 17225 | 521384 | 111722 | 134.00 | 0.00107 |
| spectra-cluster MOST |  | 1093792 | 248677 | 17183 | 525685 | 111305 | 129.38 | 0.00115 |
| spectra-cluster AVERAGE |  | 1093792 | 247808 | 17202 | 525716 | 111515 | 131.08 | 0.00113 |

PXD023047

| **MS name** |  | **No. MS** | **No. Unique Peptide** | **No. Unique Peptide**  **（FDR=1）** | **No. PSMs** | **No. PSMs**  **（FDR=1）** | **composite cluster RawScore mean** | **composite cluster Evalue mean** |
| --- | --- | --- | --- | --- | --- | --- | --- | --- |
| Original MS |  | 109333 | 23649 | 8636 | 83138 | 52617 | - | - |
| MARACluster BEST |  | 56921 | 21032 | 8053 | 43502 | 21886 | 73.33 | 0.00400 |
| MARACluster BIN |  | 63306 | 21114 | 7904 | 43565 | 21424 | 72.05 | 0.00394 |
| MARACluster MOST |  | 63306 | 21119 | 7836 | 43442 | 21095 | 66.25 | 0.00400 |
| MARACluster AVERAGE |  | 63306 | 21068 | 7856 | 43470 | 21221 | 63.60 | 0.00397 |
| spectra-cluster BEST |  | 46993 | 19915 | 7941 | 35362 | 17950 | 73.99 | 0.00361 |
| spectra-cluster BIN |  | 52262 | 19878 | 7824 | 35135 | 17633 | 71.22 | 0.00405 |
| spectra-cluster MOST |  | 52653 | 20018 | 7672 | 35226 | 17159 | 65.46 | 0.00386 |
| spectra-cluster AVERAGE |  | 52653 | 20001 | 7710 | 35308 | 17352 | 60.63 | 0.00389 |

PXD021518

| **MS name** |  | **No. MS** | **No. Unique Peptide** | **No. Unique Peptide**  **（FDR=1）** | **No. PSMs** | **No. PSMs**  **（FDR=1）** | **composite cluster RawScore mean** | **composite cluster Evalue mean** |
| --- | --- | --- | --- | --- | --- | --- | --- | --- |
| Original MS |  | 286410 | 104773 | 16427 | 172466 | 34173 | - | - |
| MARACluster BEST |  | 245917 | 102201 | 16073 | 157576 | 27290 | 79.54 | 0.00275 |
| MARACluster BIN |  | 265028 | 102170 | 15617 | 157429 | 26402 | 77.85 | 0.00263 |
| MARACluster MOST |  | 265028 | 102183 | 15575 | 157443 | 26320 | 70.48 | 0.00275 |
| MARACluster AVERAGE |  | 265028 | 102168 | 15605 | 157435 | 26381 | 71.93 | 0.00262 |
| spectra-cluster BEST |  | 241472 | 101782 | 15752 | 154257 | 25259 | 76.55 | 0.00272 |
| spectra-cluster BIN |  | 259656 | 101620 | 15363 | 153957 | 24532 | 73.29 | 0.00279 |
| spectra-cluster MOST |  | 259957 | 101666 | 15090 | 154030 | 24045 | 66.77 | 0.00271 |
| spectra-cluster AVERAGE |  | 259957 | 101630 | 15203 | 153984 | 24237 | 63.80 | 0.00262 |

PXD023361

| **MS name** |  | **No. MS** | **No. Unique Peptide** | **No. Unique Peptide**  **（FDR=1）** | **No. PSMs** | **No. PSMs**  **（FDR=1）** | **composite cluster RawScore mean** | **composite cluster Evalue mean** |
| --- | --- | --- | --- | --- | --- | --- | --- | --- |
| Original MS |  | 38286 | 16109 | 1016 | 20223 | 2588 | - | - |
| MARACluster BEST |  | 34398 | 15359 | 1002 | 18606 | 2167 | 66.64 | 0.00282 |
| MARACluster BIN |  | 36520 | 15836 | 996 | 19142 | 2168 | 70.71 | 0.00240 |
| MARACluster MOST |  | 36520 | 15773 | 997 | 19134 | 2164 | 66.61 | 0.00290 |
| MARACluster AVERAGE |  | 36520 | 15806 | 998 | 19162 | 2164 | 66.66 | 0.00269 |
| spectra-cluster BEST |  | 33696 | 15225 | 1001 | 18177 | 2034 | 66.52 | 0.00288 |
| spectra-cluster BIN |  | 35707 | 15663 | 1003 | 18678 | 2054 | 70.54 | 0.00268 |
| spectra-cluster MOST |  | 35721 | 15641 | 998 | 18708 | 2035 | 66.68 | 0.00293 |
| spectra-cluster AVERAGE |  | 35721 | 15664 | 998 | 18726 | 2039 | 66.83 | 0.00273 |

### **Supplementary Note 3: Analysis of phosphorylation sites identification of dataset PXD008355.**

| Cluster methods |  | Methods | No. PSMs  with phosphorylation sites | No. phosphorylation sites | No. PSMs with identical sites | No. PSMs with different sites | No. PSMs with different peptide | No. unique PSMs |
| --- | --- | --- | --- | --- | --- | --- | --- | --- |
| MaRaCluster |  | BEST | 66914 | 81238 | 63165 | 2683 | 57 | 1009 |
|  |  | BIN | 68429 | 83091 |  |  |  | 2524 |
|  |  | BEST | 66914 | 81238 | 62979 | 1988 | 25 | 1922 |
|  |  | MOST | 66555 | 80685 |  |  |  | 1563 |
|  |  | BEST | 66914 | 81238 | 62872 | 2626 | 39 | 1377 |
|  |  | AVERAGE | 67465 | 81924 |  |  |  | 1928 |
|  |  | BIN | 68429 | 83091 | 63180 | 2436 | 27 | 2786 |
|  |  | MOST | 66555 | 80685 |  |  |  | 912 |
|  |  | BIN | 68429 | 83091 | 64576 | 2177 | 20 | 1656 |
|  |  | AVERAGE | 67465 | 81924 |  |  |  | 692 |
|  |  | MOST | 66555 | 80685 | 62958 | 2307 | 23 | 1267 |
|  |  | AVERAGE | 67465 | 81924 |  |  |  | 2177 |
| spectra-cluster |  | BEST | 91195 | 109877 | 89161 | 1494 | 26 | 514 |
|  |  | BIN | 92202 | 111230 |  |  |  | 1521 |
|  |  | BEST | 91195 | 109877 | 89145 | 1145 | 20 | 885 |
|  |  | MOST | 91259 | 109961 |  |  |  | 949 |
|  |  | BEST | 91195 | 109877 | 89038 | 1475 | 24 | 658 |
|  |  | AVERAGE | 91729 | 110648 |  |  |  | 1192 |
|  |  | BIN | 92202 | 111230 | 89422 | 1343 | 16 | 1421 |
|  |  | MOST | 91259 | 109961 |  |  |  | 478 |
|  |  | BIN | 92202 | 111230 | 90180 | 1164 | 12 | 846 |
|  |  | AVERAGE | 91729 | 110648 |  |  |  | 373 |
|  |  | MOST | 91259 | 109961 | 89222 | 1340 | 15 | 682 |
|  |  | AVERAGE | 91729 | 110648 |  |  |  | 1152 |
